# Loss of *p16INK4a* in neuroblastoma cells induces shift to an immature state with mesenchymal characteristics and increases sensitivity to EGFR inhibitors

**DOI:** 10.1101/2021.10.27.465137

**Authors:** Nil A. Schubert, Sander R. van Hooff, Linda Schild, Kimberley Ober, Marjolein Hortensius, Kim van den Handel, Anke H.W. Essing, Bianca Koopmans, Manon Boeije, Natalie Proost, Marieke van de Ven, Selina Jansky, Sabine A. Stainczyk, Umut H. Toprak, Frank Westermann, Selma Eising, Jan J. Molenaar, Marlinde L. van den Boogaard

## Abstract

Homozygous inactivation of the *CDKN2A* locus is one of the most common genomic aberrations in human cancer. The locus codes for two unrelated and distinctly regulated proteins: p14ARF and p16INK4a, which inhibit MDM2 and CDK4/6, respectively. Loss of *CDKN2A* is also a recurrent event in relapsed neuroblastoma, a childhood tumour that arises from neural crest cells. To examine the consequences of the loss of the two distinct gene transcripts in neuroblastoma, we used the CRISPR-Cas9 system to knockout *p14, p16* and *p14*+*p16* in SY5Y cells. RNA sequencing of the transcriptome revealed a striking shift towards an immature Schwann cell precursor-like phenotype with mesenchymal characteristics, specifically in the *p16* and *p14*+*p16* knockouts. High-throughput drug screening of *p16* and *p14*+*p16* knockout clones identified a large increase in sensitivity to EGFR inhibitors. On protein level, we were able to confirm that EGFR pathway activation is higher in *p14*+*p16* knockout cells and that treatment with the EGFR inhibitor afatinib resulted in higher levels of apoptosis. Afatinib also reduced tumour growth *in vivo* in xenografts transplanted with *p14*+*p16* knockout SY5Y cells. Overall, our study suggests that *CDKN2A* deletion in neuroblastoma relates to a phenotypic shift towards a more progenitor like state and increases sensitivity to EGFR inhibitors.

## INTRODUCTION

Copy number loss of *CDKN2A* is among the most common genetic alterations in human cancers, both adult^1^ and paediatric^2^. The composition of the *CDKN2A* locus is quite unique for the human genome, as it codes for two unrelated proteins: p14ARF and p16INK4a (hereafter referred to as p14 and p16). The two genes share exon 2 and 3 but have distinct first exons, start sites and promoters, resulting in different open reading frames. Both proteins play a key role in prominent cancer pathways regulating cell cycle control. Although inactivation of only one gene can occur (usually via point mutation or promoter methylation) the typical cancer-related mutation affects both alleles and thus both pathways^3^. Expression of p14, which serves as an activator of p53, is typically low but enhanced upon oncogenic stimulation. The protein binds to and sequesters the E3 ubiquitin ligase MDM2 to the nucleolus, thereby indirectly stabilizing p53, which can then exert its function as a transcription factor for genes involved in apoptosis and cell cycle arrest. The pRb pathway is regulated by p16, which inhibits the CDK4/6-cyclin D1 complex, thereby preventing the phosphorylation of pRb. E2F transcription factors retain bound to pRb and expression of S-phase entry genes is not initiated. In other words, loss of *CDKN2A* results in increased MDM2 and CDK4/6 activity and hence disturbed p53 and pRb signalling. Negative feedback loops exist between pRb and p16^4^, as well as between p53 and p14^5^. Moreover, crosstalk between the two pathways has also been described, e.g., via the cyclin-dependent kinase inhibitor (CKI) p21, which is a target gene of p53^6^. Thus, *CDKN2A* copy number losses may contribute to rapid cell cycle progression and decreased rates of apoptosis.

Loss of *CDKN2A* has also been identified in neuroblastoma^7,8^. Neuroblastoma is the most common extracranial paediatric solid tumour^9^. These tumours arise from neural crest cells and are typically located in the adrenal medulla or along the sympathetic chain^10,11^. Upon refractory or relapsed disease, the average overall survival of 50-60% decreases to less than 10%^12–14^. Despite a paucity of somatic mutations in neuroblastoma, we and others previously showed that there is a general enrichment of mutations in refractory or relapsed tumours^15,16^. While homozygous *CDKN2A* deletions occur in ∼3% of primary neuroblastomas^7,8,17–19^, our group previously identified this copy number loss in 13% of relapse tumours but not in the matching primary samples. Carr-Wilkinson et al. also reported a higher frequency of p53/ MDM2/p14 pathway abnormalities in relapsed neuroblastoma^19^.

*CDKN2A* loss is considered an actionable event, meaning that the inhibition of downstream proteins is thought to interfere with uncontrolled cell division. However, in a previous study, we have shown no significant change in sensitivity to MDM2 and CDK4/6 inhibitors in two neuroblastoma cell lines with homozygous loss of *CDKN2A*^20^. We designed this study to address the consequences of *p14* inactivation, *p16* inactivation and the combined loss in neuroblastoma. While several studies examined the knockdown or knockout of either *p14* (or *p19* in mice)^21,22^ or *p16*^*23*,24^, we are unaware of studies comparing these two situations. To study these individual effects, we generated different knockouts in SY5Y neuroblastoma cells using CRISPR-Cas9. We performed RNA sequencing on clonal cell lines with *p14* and/or *p16* knockout to gain more insight in gene expression changes, which revealed a clear shift towards a more progenitor-like, mesenchymal state upon knockout of *p16* but not *p14*. Subsequent high-throughput drug screening revealed a striking increase in EGFR inhibitor sensitivity of the *p16* knockout cells. Afatinib also reduced tumour growth in *p16* knockout xenografts.

## MATERIAL & METHODS

### Cell culture

All cell lines were cultured in Dulbecco’s Modified Eagle Medium (DMEM), supplemented with 10% fetal bovine serum, 2 mM L-glutamine, 1% non-essential amino acids, 100 U/mL penicillin and 100 mg/mL streptomycin. Cells were grown at 37 °C and 5% CO2 and routinely tested for mycoplasma infection.

### Establishments of knockout lines (gRNA cloning, virus production, clone selection)

Knockout of *p14ARF* was obtained by targeting *CDKN2A’s* exon 1β, using the single guide RNA (sgRNA) CACCGGCGGCGAGAACATGGTGCGC (“C2”) or CACCGGCACGCGCGCCGAATCCGGA (“C3”). By targeting exon 1α with sgRNA CACCGACGCACCGAATAGTTACGGT (“C13”) *p16INK4a* was knocked out, and knockouts of CDKN2A (*p14*+*p16*) were established by targeting exon 2 with sgRNA CACCGGTGGCGGGGTCGGCGCAGTT (“C7”) or CACCGGACCCGTGCACGACGCTGCC (“C8”). The sgRNAs were designed using the previously available Zhang lab CRISPR Design Tool. sgRNAs were cloned into the lentiCRISPRv2 plasmid (a gift from Feng Zhang, Addgene plasmid #52961) as described by the Zhang lab^25,26^. For lentivirus production, the obtained plasmids, in combination with pMD2G, pRRE and pSRV/REV packaging vectors, were transfected in HEK293T cells using FuGENE® (Promega). DMEM medium, containing the virus, was collected, filtered and then concentrated by ultracentrifugation. SY5Y neuroblastoma cells were subsequently transduced with 10 μL of the concentrated virus and selected for successful transduction with puromycin. Cells were then plated at low densities to ensure clonal outgrowth of single cells, after which several colonies were picked, expanded, and checked for the correct knockout using Sanger sequencing and Western blot analysis.

### DNA validation of knockout cells

DNA was isolated using the Wizard® SV Genomic DNA Purification System kit (Promega). The DNA was amplified with a polymerase chain reaction (PCR), using primers for exon 1β/*p14ARF* (F: AGCAATGAGATGACCTCGCTT, R: CACCAAACAAAACAAGTGCCG), exon 1α/*p16INK4a* (F: TGACACCAAACACCCCGATT, R: AGAATCGAAGCGCTACCTGA) or exon 2/*CDKN2A* (F: TTACCACATTCTGCGCTTGG or *TTAGACACCTGGGGCTTGTG*, R: TGGAAGCTCTCAGGGTACAAAT or ATGGTTACTGCCTCTGGT). Clone C13.16 was validated using the CloneJET PCR Cloning Kit (Thermo Scientific) and the primers for exon 1α/*p16INK4a*, according to the manufacturer’s protocol. Purified PCR products were sent to Macrogen for Sanger sequencing using the same primers. Results were analyzed using the SnapGene software and the online tool TIDE^27^.

### Western blot analysis

Cells were grown in 10 cm dishes and, if needed, treated with different drug concentrations or DMSO for 24 or 48 hours. Cells were washed with phosphate-buffered saline (PBS) and then harvested and lysed with Laemmli buffer (20% Glycerol, 4% sodium dodecyl sulphate and 100mM Tris HCl), supplemented with 0.1% NAF, 0.1% Na_2_VO_3_ and cOmplete^™^ mini protease inhibitor cocktail (Roche). Protein content was measured using the Bio-Rad DC protein assay. Equal amounts of protein were loaded and run on Bio-Rad Mini-Protean^®^ TGX™ 10% gels. The Trans-Blot^®^ Turbo™ Transfer System (Bio-Rad) was used to transfer proteins onto a polyvinylidene difluoride (PVDF) membrane. Membranes were blocked in ECL advance blocking reagent (GE Healthcare) in TBS-Tween 0.1% and incubated with primary antibodies overnight at 4 °C. Incubation with secondary antibodies was done at room temperature for one hour. Protein bands were visualised using either the Chemidoc^™^ Touch (Bio-Rad) or the Odyssey CLx (LI-COR).

### Antibodies

Primary antibodies were p16INK4a (AF5779, R&D Systems, 1:200), MDM2 (A2716, Calbiochem), CDK4 (AHZ0202, Invitrogen), EGFR (4267, Cell Signaling), p-EGFR (3777, Cell Signaling), AKT (4691S, Cell Signaling), p-AKT (4056S, Cell Signaling), MEK (8727S, Cell Signaling), p-MEK (9154S, Cell Signaling), PARP (95425, Cell Signaling) and β-actin (ab6276, Abcam). All primary antibodies were diluted 1:1000 with 2% ABA blocking buffer (Amersham, #RPN418V) in TBST, unless stated otherwise. Secondary antibodies for chemiluminescence were anti-mouse (NXA931V, GE Healthcare), anti-rabbit (NA9340V, GE Healthcare), anti-goat (KO615, Santa Cruz) and for fluorescence anti-mouse (926-32210, LI-COR) and anti-rabbit (926-32211, LI-COR). All secondary antibodies were diluted 1:5000.

### RNA sequencing and establishment of signatures

Samples were harvested when cells were seeded for the high-throughput screen. A duplicate sample for wildtype SY5Y cells was harvested at a later time point. Cells were spun down, resuspended in TRIzol^™^ (Invitrogen) and chloroform was added to separate the phases. RNA was precipitated by adding 100% RNA-free ethanol to the aqueous phase. Finally, RNA was extracted using the RNeasy Mini Kit (Qiagen). Illumina sequencing libraries were prepared using the Truseq RNA stranded polyA kit (Illumina) and sequenced with 2×50p paired-end sequencing on an Illumina NextSeq2000 system. After sequencing the data was aligned to genome build GRCh37.p13 and reads were summarized per gene with the R package Rsubread^28^ using Ensembl gene build GRCh37.v74. Counts per gene were normalized to TPM values (Transcripts Per Kilobase Million) and log transformed and corrected for experimental batch using ComBat as implemented in the R package sva^29^. Genes differentially expressed against the wildtype cell line were identified using the DESeq2^30^ R package taking along the experimental batch as a variable in the design. GSEA analysis was performed using the R package fgsea with the hallmark gene sets from MsigDB v7.2^31,32^. The p16 signature was composed of the shrunken log fold changes of the 250 most significantly regulated genes in the *p16* and *p14*+*p16* knockout cell lines (as a single group). No batch correction was applied in this procedure. The signature contained 118 upregulated and 132 downregulated genes. Scores for the expression of the p16 up and p16 down gene signatures in single developing adrenal medullary cells were determined using the AddModuleScore function in R package Seurat v.3.1^33^ with 100 control genes per analyzed gene. Briefly, the average expression of the gene signature was determined in individual cells and subtracted by the expression of the control gene sets. Expression scores were visualized on the UMAP embedding calculated for normal developing adrenal medullary cells published by Jansky et al^34^.

### High-throughput compound screen

Screening experiments and processing were performed by the high-throughput screening facility of the Princess Máxima Center (https://research.prinsesmaximacentrum.nl/en/core-facilities/high-throughput-screening). Cells were seeded in 384-well microplates using the Multidrop™ Combi Reagent Dispenser (Thermo Scientific). Seeding densities (cells/well) were as follows: 3000 SY5Y, 15000 SY5Y-C2.1, 6000 SY5Y-C3.6, 2000 SY5Y-C13.7, 5000 SY5Y-C13.16, 3000 SY5Y-C7.3 and 2500 SY5Y-C8.10. After 24 hours, the 197 library compounds dissolved in DMSO or MilliQ (Supplementary Table 1) were added in duplicate using the Echo 550 dispenser (Beckman Coulter). Final drug concentrations ranged standard between 0.1 nM and 10 μM, although some drugs were, if necessary, added on lower and higher concentrations. Control samples were treated with the appropriate concentrations of solvents (0.25% of DMSO or Milli-Q). After 72 hours of treatment, cell viability was measured using the 3-(4,5-dimethylthiazol-2yl)-2,5-diphenyltetrazolium (MTT) assay^35^ and compared to DMSO-treated wells as positive controls and staurosporine-treated (10 μM) wells as negative control. Data were normalised to DMSO-treated cells (defined as 100% viability) and empty controls (0% viability). Subsequently, IC_50_ values (half maximal concentration that inhibits viability) were calculated by determining the concentrations of the drug needed to achieve a 50% reduction in cell viability, using the extension package drc in the statistic environment of R studio (version 4.0.2). Quality of the screens was approved after assessment of cell growth (absorbance signal of t72/t0), negative, positive and empty controls and the amount of variability between duplicates.

### Cell viability assay (compound validation)

Cells were seeded in quadruplicates in 384-wells plates at densities of 2000-15,000 cells per well, depending on the cell line, see above. Inhibitors were added after 24 hours using a five-fold concentration range from 0.128 nM to 50 μM with the D300e Digital Dispenser (TECAN). After 72 hours, cell viabilities were measured using the MTT assay and compared to solvent-treated controls. Two replicate experiments were performed.

### Inhibitors

EGFR inhibitors erlotinib (HY-50896), afatinib (HY-10261) and sapitinib (HY-13050) and ALK inhibitor lorlatinib (HY-12215) were all purchased from MedChemExpress. The ALK inhibitor alectinib (S2762) was obtained from SelleckChem.

### Cell cycle analysis

Cells were seeded in 6 cm dishes and treated with afatinib (50, 500 and 5000 nM) or DMSO after 24 hours. After 48 hours on treatment, attached and floating cells were harvested, spun down and washed with PBS. Subsequently, cells were washed twice with PBS with 4 mM ethylenediaminetetraacetic acid (EDTA) and spun down. Cells were then resuspended until single cells were obtained and counted. One million cells were stained with Vybrant® DyeCycle^™^ Violet Stain (Invitrogen) with a final concentration of 5 μM. Cells were then incubated in the dark at 37 °C for 30 minutes. Subsequently, DNA content was measured using the CytoFLEX LX Flow Cytometer (Beckman Coulter) and analyzed using the accompanying CytExpert software. Three replicate experiments were performed.

### Xenograft experiments

2.5×10^6^ cells of SY5Y-C7.3 or SY5Y-C8.10 were xenografted into a flank of 6-8-week-old NMRI nu^-^/nu^-^ mice (obtained from Janvier Labs). Three times a week, body weight and tumour size were measured; latter was measured using a caliper and tumour volume (mm^3^) was determined with the formula ½(length * width^2^), with length representing the largest tumour diameter and width the perpendicular diameter. Treatment was initiated once tumours reached a size of 250 mm^3^. Mice (n = 6-8 per group) were treated with afatinib (15 mg/kg) or vehicle (0,5% methylcellulose / 0,4% Tween80) orally for 21 consecutive days and sacrificed once tumour volumes exceeded 1500 mm^3^. Animals for pharmacodynamic measurements (n = 3 per group) were sacrificed after 5 days of treatment. Ethical approval was obtained under project number AVD30100202011584 (EGP no. 10064).

## RESULTS

### Generation of *p14, p16* and *p14*+*p16* knockout SY5Y cells

To study the effects of *p14* and *p16* knockout, we selected the commonly used neuroblastoma cell line SY5Y. Like the Shep2 cell line, SY5Y is a subclone derived from the SKNSH cell line^36^. While Shep2 cells carry a homozygous deletion of *CDKN2A*, SY5Y cells have two wildtype copies of *CDKN2A*. We, therefore, hypothesised that SY5Y cells have the proper genomic background for mimicking the loss of the *CDKN2A* locus. To create the distinct knockouts, we cloned guide RNAs (gRNAs) specifically targeting *p14, p16* or *p14*+*p16* loci (Fig. **1a**) into the pLentiCRISPRv2 plasmid and transduced wildtype SY5Y cells. After selection and clonal expansion of single cells, *p14* and *p16* knockout in the clones was confirmed using Sanger sequencing, in combination with the online tool TIDE^27^, and Western blot analysis (Fig. **1b**). For each target, we selected two clones to proceed with. Compared to the reference sequence, clear out-of-frame interruptions were seen in all sequences (Fig. **1c**). In concordance, p16 protein was absent in all *p16* knockout clones (C13.7, C13.16, C7.8, and C8.10) (Fig. **1d**). Unfortunately, the detection of wildtype p14 protein proved to be complicated due to non-specific antibodies, so we initially had to rely on DNA validation for these knockouts.

**Figure 1.**
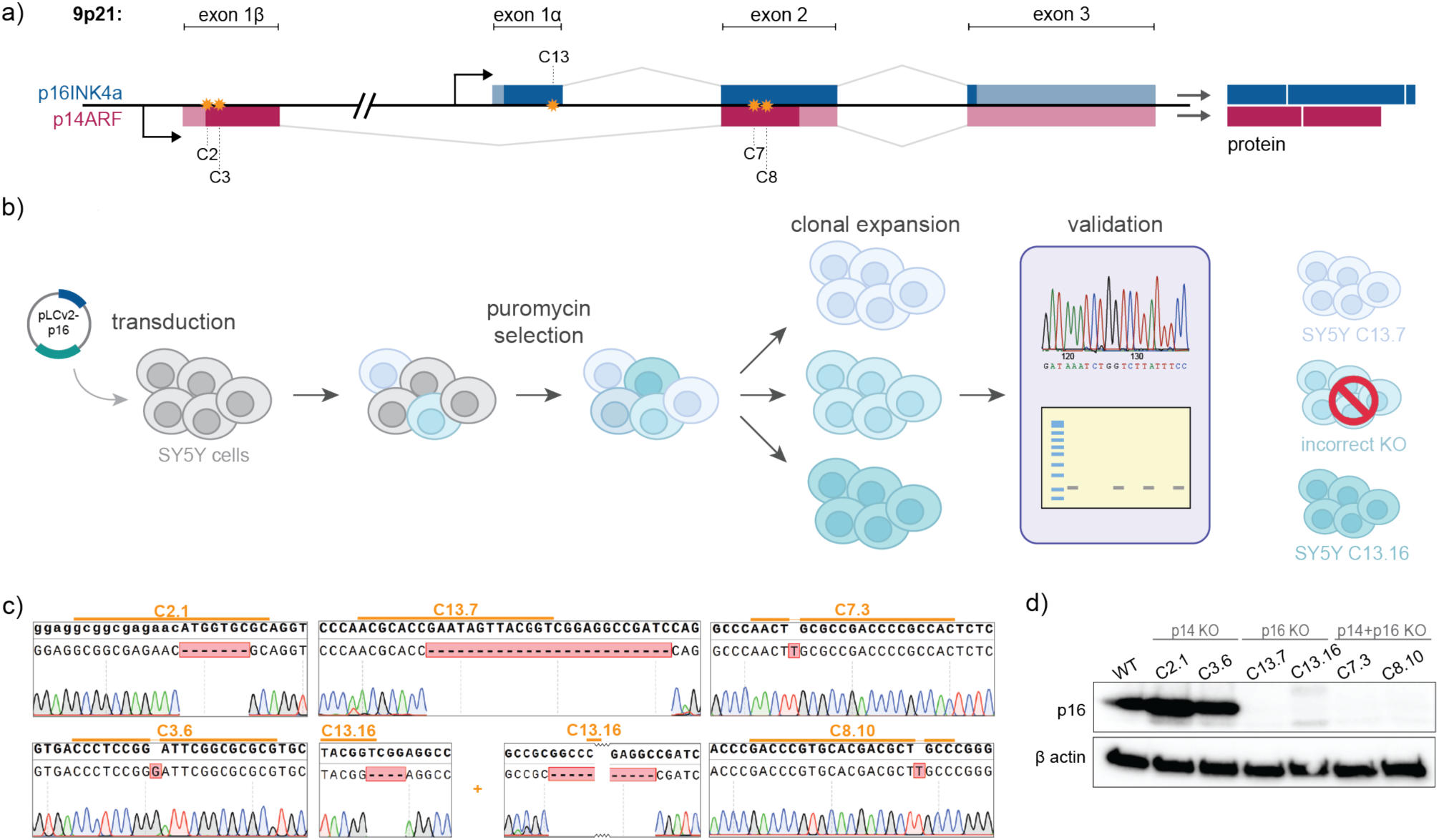
Knockouts of *p14, p16* and *p14*+*p16* were generated in SY5Y neuroblastoma cells. **a)** Structure of the *CDKN2A* locus and genomic locations targeted by the different gRNAs. gRNAs C2 and C3 were directed towards exon 1β and thus *p14*, while C13 targeted exon 1α (*p16*) and C7 and C8 exon 2 (*p14*+*p16*). **b)** Schematic representation of the establishment of clonal cell lines. SY5Y cells were transduced with the different gRNA-containing plasmids and selected with puromycin. Seeding the cells at very low densities ensured clonal expansion of single cells. Knockouts were validated using Sanger sequencing and Western blot. Two knockout clones were selected for each target. **c)** Sequences of the clonal cell lines that were selected for this study. All cell lines presented with an out-of-frame insertion or deletion. C13.16 had a four base pair deletion on one allele and a deletion of 83 base pairs on the other. **d)** Protein levels of p16 in all selected clones. Protein is indeed absent in cells with a knockout of *p16*.

### Loss of p16 has a significant effect on the transcriptome

To investigate the effects of *p14* and/or *p16* loss on the transcriptome, we performed RNA sequencing on all six clonal cell lines and two replicates of the wildtype SY5Y cells. Visualisation of the RNA reads showed a clear gap or mutation at the location of the gRNAs, corresponding in length to the deletion or insertion identified with Sanger sequencing. Thus, we concluded that we successfully knocked out *p14*. Subsequent differential gene expression analysis of knockout clones compared to the wildtype (p-value of <0.05 and a log fold change of >1), showed a minor effect of *p14* knockout, while *p16* and *p14*+*p16* knockout clones showed striking changes in the gene expression profile (Fig. **2a**). Additional clustering based on a Pearson correlation of the 500 genes with the most variable expression, showed that *p14* knockouts clustered together with the wildtype cells, whereas *p16* and *p14*+*p16* knockouts form a distinct cluster (Fig. **2b**). This observation is confirmed if we cluster the top 50 differentially expressed genes in the p*14*+*p16* knockouts (Fig. **2c**) and *p16* knockouts compared to the *p14* knockouts (Supplementary Fig. **1**).

**Figure 2.**
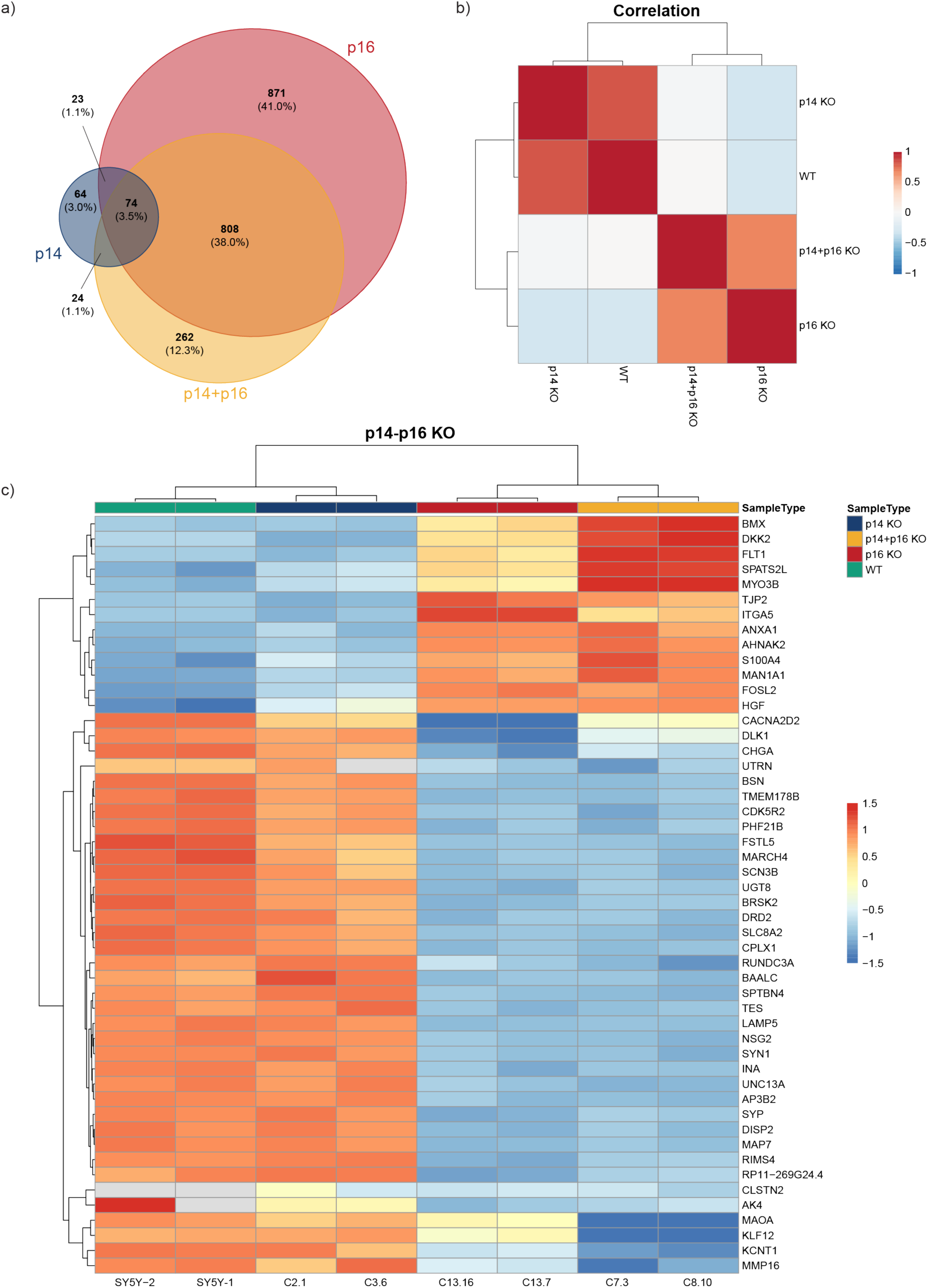
Gene expression of the different knockout cell lines. **a)** Venn diagram showing differentially expressed genes per knockout compared to wildtype. Genes were selected using an adjusted p-value of <0.05 and an absolute log fold change >1. **b)** Correlation of the average expression between the different knockouts. The 500 most variable genes were used to calculate the Pearson correlation coefficients. **c)** Heatmap showing the expression (transcripts per million) of the 50 most differentially expressed genes (*p14*+*p16* knockout versus wildtype) with an absolute log fold change >1. Genes are standardised row-wise.

### Loss of *p16* induces a shift towards a more immature, mesenchymal state

To study the phenotypic switch in *p16* knockout clones in more detail, we performed gene set enrichment analysis (GSEA) using the Hallmark gene sets of the Molecular Signatures Database (MSigDB)^32^. This analysis indicated significant overlap in the top 20 gene sets that were altered upon *p16* and *p14*+*p16* knockout, especially in the upregulated gene sets (Fig. **3a+b**). The combined top hit was the upregulation of genes linked to epithelial to mesenchymal transition (EMT), suggesting a switch to a more mesenchymal phenotype after *p16* knockout (Fig. **3c**). Jansky et al. recently published the developmental trajectory of neuroblastoma based on single-nucleus RNA sequencing^34^. To see what the effect of the *p16* knockout was when viewed in the context of this developmental trajectory, we generated *p16* knockout upregulated (p16 KO up) and downregulated (p16 KO down) gene expression signatures and determined their expression scores in the individual developing adrenal medullary cells. The genes from the p16 up signature are highly expressed in Schwann cell precursor (SCP) cells, whereas the genes from the p16 down signature are associated with high expression in the more differentiated neuroblasts, connecting progenitor cells and chromaffin cells (Fig. **3d**). Overall, these results indicate that *p16* knockout induces a strong transcriptomic switch into a more immature SCP-like phenotype with mesenchymal characteristics.

**Figure 3.**
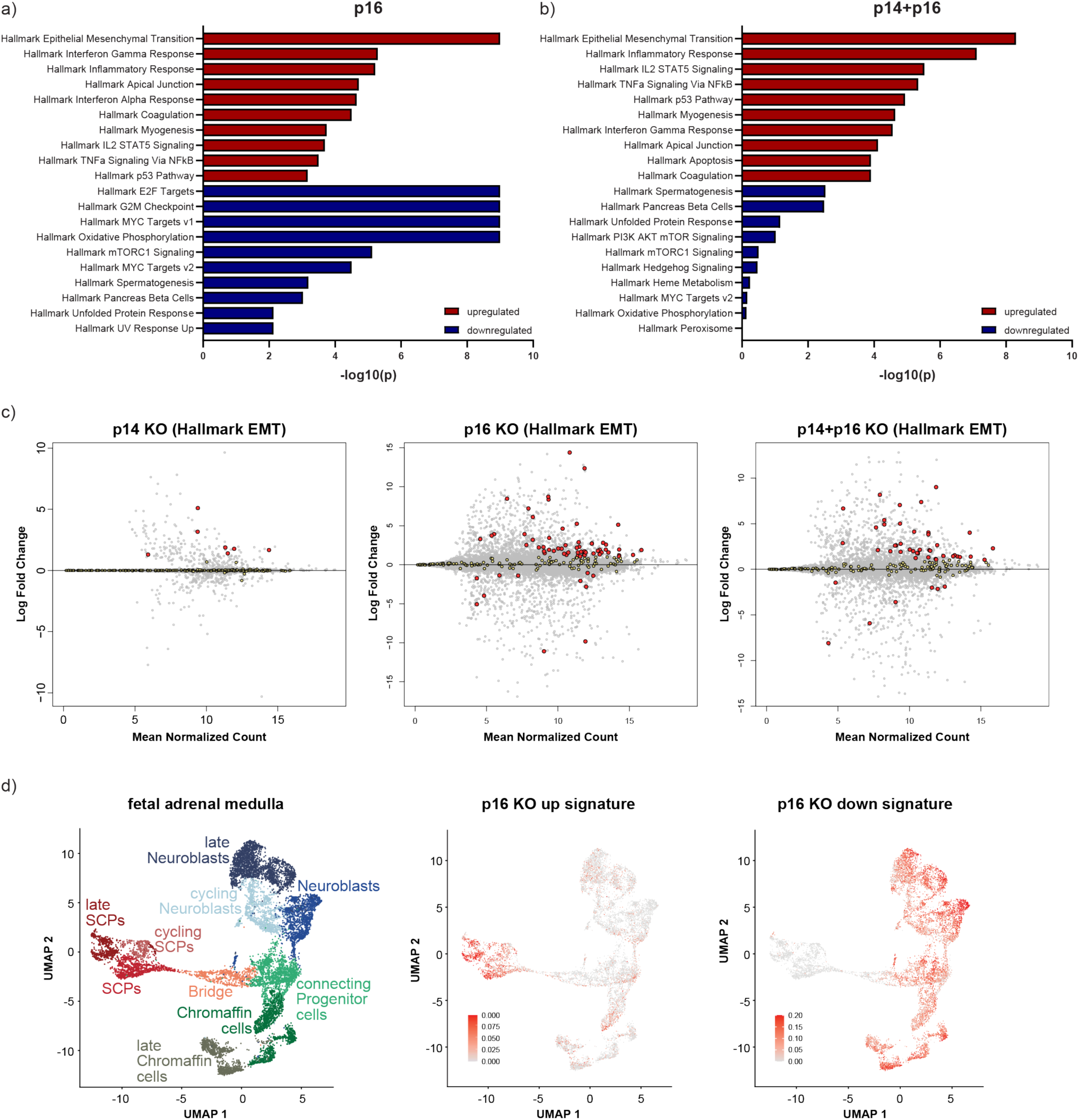
Gene set enrichment analyses and signatures of the knockout cell lines. Top 10 most upregulated and downregulated gene sets in the *p16* **(a)** and *p14*+*p16* **(b)** knockout cell lines. Gene sets are ranked based on the adjusted p-value. **c)** MA plots marking the individual genes of the Hallmark Epithelial Mesenchymal Transition gene set. The x-axis shows the log2 of mean normalised counts, the y-axis the shrunken log fold change of the different knockout cell lines versus the wildtype cell line. Red dots indicate gene set members with an absolute log fold change >1, the yellow dots those with an absolute log fold change <1. In grey all other genes are shown. **d)** Diffusion map of adrenal medullary cells colored by chromaffin cell lineage pseudotime trajectory (adjusted with permission from Jansky et al.^34^) (left) and expression of genes from the p16 KO up (middle) and p16 KO down (right) signatures in UMAP embedding of the adrenal medullary cells. The log-normalised gene expression is indicated by color (middle and right).

### Loss of *p16* increases sensitivity to EGFR inhibitors

Since such a significant shift in gene expression and phenotype could very well result in altered sensitivity to targeted compounds, we subjected all knockout cell lines and the SY5Y parental line to a 72-hour high-throughput compound sensitivity screen using a library of 197 drugs (Fig. **4a**). These drugs are either currently used in cancer treatment or clinical trials or are far along in (pre)clinical development (Table **1**). Cell viability curves were established for each drug in each cell line, and the half maximal concentration that inhibits the viability (IC_50_) and area under the curve (AUC) values were calculated. Comparing all AUC values for the three types of knockout lines with wildtype SY5Y, showed that single knockout of *p14* had only a minor effect on drug sensitivity (Fig. **4b**). More significant shifts in compound sensitivity were seen in the *p16* and the *p14*+*p16* knockout cells. We subsequently divided the cell lines in two groups: the *p16* wildtype group (SY5Y wildtype and *p14* knockout cells) and the *p16* knockout group (*p16* and *p14*+*p16* knockout cells). Analysis of differential compound sensitivities showed a highly significant increase in sensitivity for all nine EGFR inhibitors included in the library (p<0.01 for 8/9 inhibitors; p<0.05 for the remaining inhibitor) (Fig. **4c+d**). Additionally, we observed a highly significant increase in resistance for three of the nine ALK inhibitors (p<0.01) (Supplementary Fig. **2a**), which was confirmed by treating the two *p14*+*p16* knockout cell lines and wildtype cells with alectinib and lorlatinib (Supplementary Fig. **2b**). Interestingly, ALK expression was significantly lower in the *p16* and the *p14*+*p16* knockouts (Supplementary Fig. **2c**). We also identified increased resistance to BCL2 inhibitors and greater sensitivity to MEK inhibitors in these cells (Supplementary Fig. **2d+e**).

**Figure 4.**
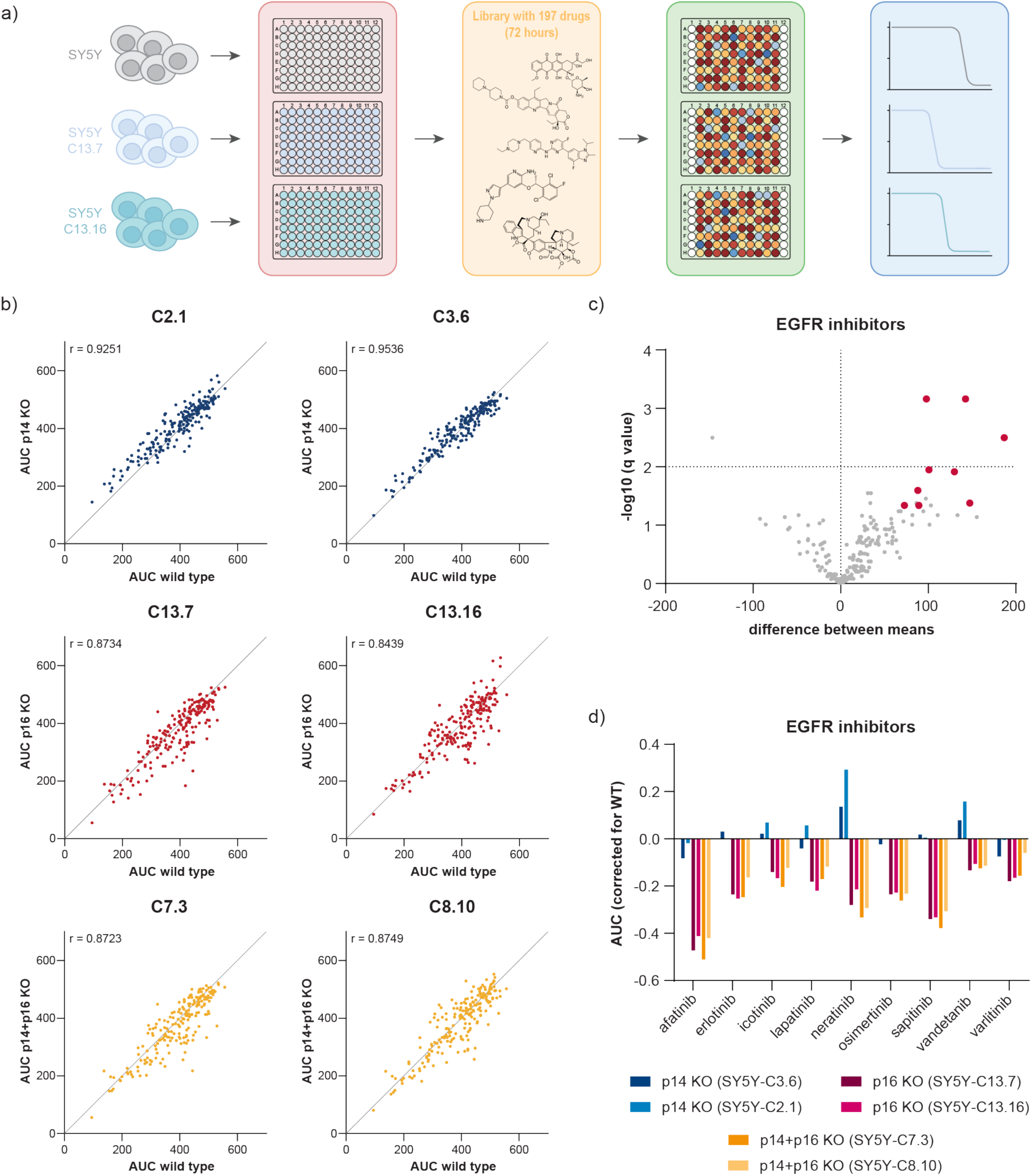
High-throughput drug screening of the different clonal cell lines. **a)** Schematic overview of the procedure. Clonal cell lines were seeded in 384-wells plates and subsequently treated with a library containing 197 drugs for 72 hours. Drug sensitivities of the knockout cell lines were compared to the sensitivities of SY5Y wildtype cells. **b)** Correlation plots for each clonal cell line, showing the area under the curve (AUC) for each drug in the knockout cell line and SY5Y wildtype cells. **c)** Volcano plot of *p16* knockout vs. *p16* wildtype AUC values. All nine EGFR inhibitors included in this experiment are highlighted in red. **d)** Waterfall plot showing the AUC values of the different knockout cell lines compared to the SY5Y wildtype cells for each EGFR inhibitor included.

In our previous study, we showed that *CDKN2A*-deleted cell lines do not respond better to MDM2 or CDK4/6 inhibition^20^. In line with these findings, our current compound sensitivity screening revealed that none of the knockouts significantly influences sensitivity to the four MDM2 and the three CDK4/6 inhibitors from the library (Supplementary Fig. **2f+g**).

### *CDKN2A* knockout increases dependency on the EGFR pathway

Because of the striking enrichment of EGFR inhibitors in our library screen, we decided to focus on the validation of this class of drugs. For the validation using a bigger concentration range, we chose afatinib, since this compound showed the greatest shift in the high-throughput screen, and erlotinib because Phase I study results in paediatric patients are already available^37^. These results confirmed that SY5Y cells with a knockout that includes *p16* are indeed more sensitive to these EGFR inhibitors, with IC_50_ values for afatinib being much lower compared to SY5Y wildtype (Fig. **5a+b**). Since the *p14*+*p16* knockouts (C7.3 and C8.10) are clinically most relevant, we decided to further focus on these clonal cell lines. Protein analysis revealed higher basal levels of both unphosphorylated and phosphorylated EGFR (p-EGFR) in the knockout cell lines (Fig. **5c**). Incremental concentrations of afatinib (for 24 hours) resulted in decreased levels of p-EGFR, as well as of the downstream proteins p-AKT and p-MEK. This decrease in pathway activation was best visible in C7.3 cells. After 48 hours of treatment, PARP cleavage already occurred at a dose of 50 nM in the knockout cells, whereas the first sign of PARP cleavage in the wildtype cells was seen with 5000 nM afatinib. Although afatinib induced cell death (indicated by the sub G1 fraction) in all cell lines, this effect was much stronger when *CDKN2A* was knocked out (Fig. **5d**). These results hint towards higher basal activity of the EGFR pathway in a *CDKN2A*-deleted background, which causes increased dependency of these cells on this pathway.

**Figure 5.**
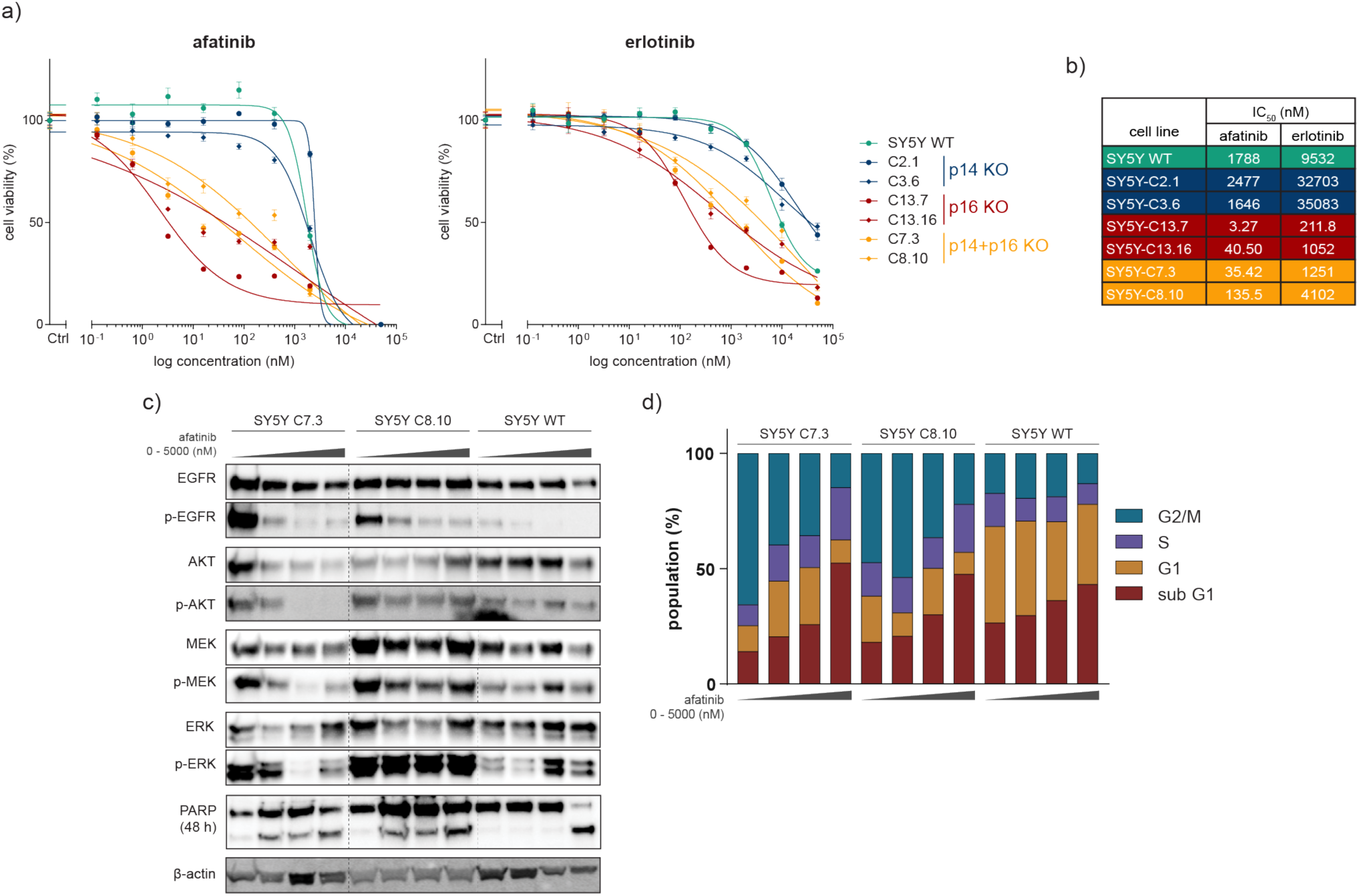
Validation of EGFR inhibitors in *CDKN2A* knockout cell lines. Survival curves of the different clonal cell lines after 72-hour treatment with afatinib and erlotinib **(a)** and the corresponding IC_50_ values **(b)**. Error bars indicate the SEM. **c)** Protein levels after treatment with increasing doses of afatinib of the two *p14*+*p16* knockout and the wildtype cell lines. PARP protein samples were harvested after 48 hours of treatment, while the rest was harvested after 24 hours. β-actin levels were comparable for all samples. Cell cycle distribution in the two *p14*+*p16* knockout and the wildtype cell lines after 48-hour treatment with increasing doses of afatinib. Afatinib concentrations used for the Western blot and FACS were 0, 50, 500 and 5000 nM.

### Afatinib reduces tumour growth in xenografts with *CDKN2A* knockout

To study the sensitivity of CDKN2A knockouts to EGFR inhibition *in vivo*, we established two xenograft models by injecting SY5Y-C7.3 and SY5Y-C8.10 cells subcutaneously into NMRI nu/nu mice. Once tumours reached a size of 250 mm^3^, the 21-day oral treatment with afatinib was started. We initially started treatment at 20 mg/kg but lowered the dose once we noticed that some mice lost a significant amount of weight. Two animals in the SY5Y-C7.3 group received two doses of 20 mg/kg before changing to 15 mg/kg. However, this change in dose did not seem to alter the response, since one of the tumours progressed despite the higher dose. Despite the lower dose, some weight loss was still observed in two C7.3 animals. After 21 days, six out of eight vehicle-treated C7.3 mice and six of seven C8.10 mice had already been sacrificed due to the size of the tumours. When end of treatment tumour volumes (either after completing 21 days or because the tumour reached a volume of 1500 mm^3^) were compared, afatinib treatment led to a significant reduction in tumour size of 61.2% compared to the vehicle control in C7.3 animals (average tumour growth of 123.0% versus 450.5%, p = 0.005) (Fig. **6a+b**). In C8.10-bearing mice, the tumours were 35.9% smaller than the vehicle controls (226.7% versus 404.7%, p = 0.0426). Median survival was extended by 16.5 and 15 days for C7.3 and C8.10, respectively (Fig. **6c**).

**Figure 6.**
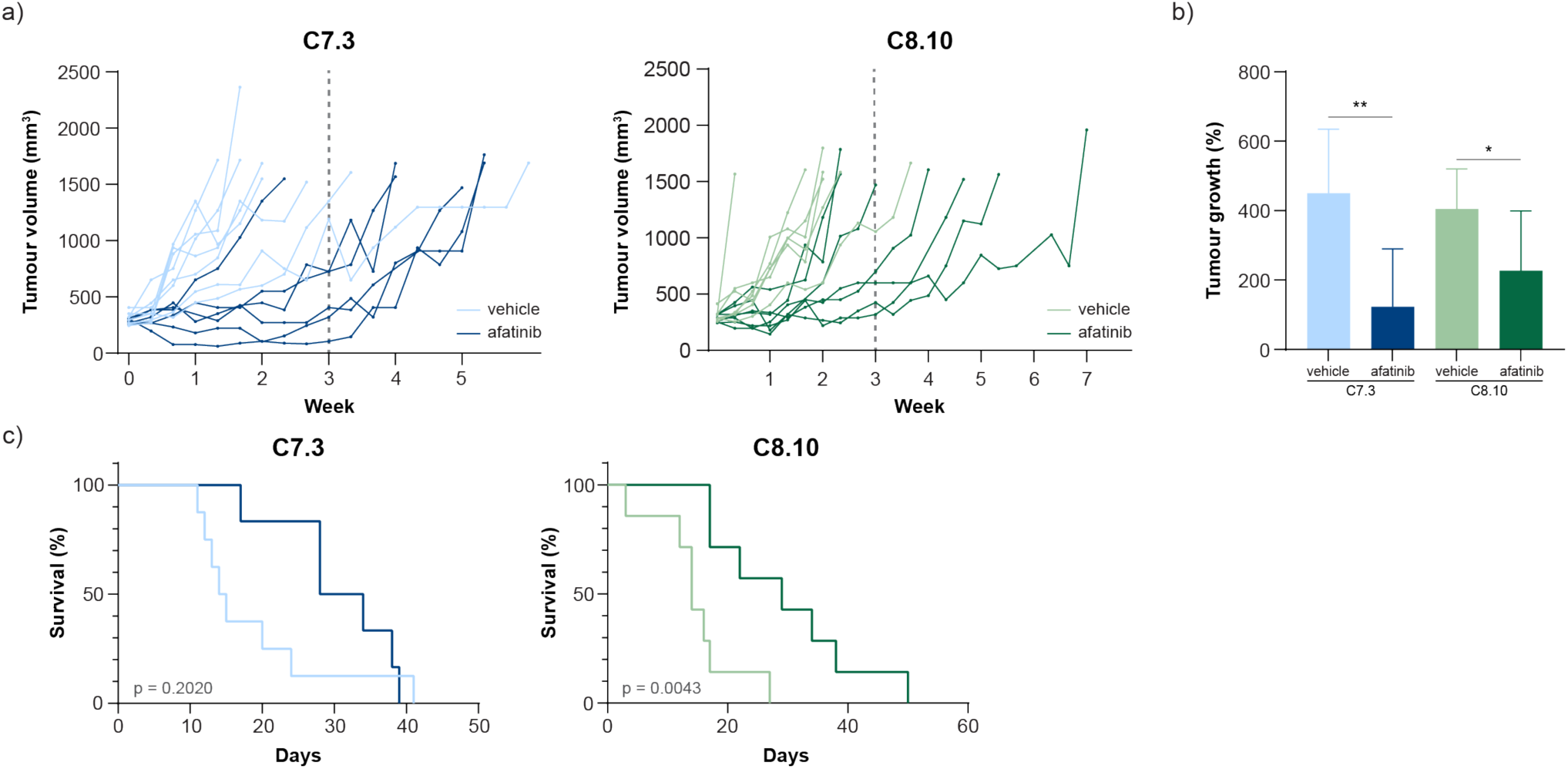
In vivo testing of afatinib in C7.3 and C8.10 (*p14*+*p16* knockout) xenografts. **a)** Tumour volume of all animals treated with either afanitib or the appropriate vehicle. Dashed line indicates the end of treatment. **b)** Average tumour growth per group. * p≤0.05; ** p≤0.01 (student’s t-test). Error bars indicate the SD. **c)** Kaplan-Meier curve showing the survival of each group. P-values were calculated using a log-rank test.

## DISCUSSION

With *CDKN2A* being one of the most frequently deleted loci in both adult and childhood cancer, it is pivotal to understand the effects of this loss better, as well as finding ways to interfere with its consequences. Here, we showed that the inactivation of *p16* has a much bigger effect on SY5Y neuroblastoma cells than *p14* inactivation. Cells shifted towards a mesenchymal phenotype upon knockout of either *p16* alone or *p14*+*p16* (*CDKN2A*) and the transcriptome reflected that of less-differentiated SCP cells. Moreover, the high-throughput drug screen revealed a changed drug response profile, of which the increased sensitivity to EGFR inhibitors was the most significant. The EGFR inhibitor afatinib induced cell death and cell cycle arrest at greater levels and lower dosages in *CDKN2A*-inactivated cells and reduced tumour growth significantly in two mouse models.

In this study, we showed that the knockout of *p14* did not lead to any striking differences in phenotype, gene expression, nor compound sensitivity in SY5Y neuroblastoma cells. It is yet unclear, why the effect of *p14* inactivation is comparably small. Dreidax et al. reported that p14 expression is epigenetically repressed and promoter activity is relatively low in SY5Y cells^8^. SY5Y cells may not depend much on the basal expression of this protein and therefore inactivation may not lead to major changes in gene expression.

Contrary to *p14* inactivation, *p16* knockout (with or without simultaneous *p14* knockout) changed gene expression exceedingly compared to wildtype SY5Y cells. All *p16* and *p14*+*p16* knockout cells switched to a more immature, mesenchymal state and became sensitive to EGFR inhibition. The transition to a mesenchymal state, typically epithelial-to-mesenchymal transition (EMT), is a common feature in cancers, enabling tumours to grow invasively and metastasise to distinct sites^38^. Mesenchymal cells are characterised by a more stem cell-like or dedifferentiated state^39^, which is in line with our finding that *p16* knockout cells resemble the more undifferentiated SCPs. Recently, Gartlgruber et al. published the mesenchymal signature activity score of 23 neuroblastoma cell lines^40^. Looking at these scores, we noticed that all mesenchymal cell lines have a heterozygous or homozygous inactivation of the *CDKN2A* locus, which also hints towards a potential correlation between a mesenchymal state and an impaired *CDKN2A* locus. It was previously shown that p16 represses EMT via miR-141 and miR-146b-5p^41^, so loss of *p16* may indeed enhance the cell’s ability to transition to a mesenchymal state. In addition, FOSL2 and/or HGF, both strongly upregulated in the *p16* and *p14*+*p16* knockout clones (Figure **2c**), might induce or contribute to the mesenchymal switch. FOSL2 is a core regulatory circuitry transcription factor in neuroblastoma, shown to regulate the mesenchymal phenotype^40^, and binding of HGF to c-Met, which regulates EMT, was shown to induce mesenchymal features in neuroblastoma^42^.

EMT is characterised by the downregulation of E-cadherin, which is typically a result of repression by the transcription factors SNAI1 and TWIST1. EGFR signalling was shown to promote EMT via the upregulation of these transcription factors^43,44^ and by stimulating reorganisation of the actin cytoskeleton in favour of EMT^45^. Thus, in our *p16* knockout cells, the mesenchymal switch could be a direct result of enhanced EGFR signalling. The question remains, however, what mechanism is behind the increased dependence on this signalling pathway upon *p16* knockout. While different (feedback) mechanisms have been identified that aid in the downregulation of EGFR via receptor endocytosis or dephosphorylation^46,47^, we are unaware of any mechanisms linking p16 directly to EGFR expression or activation in neuroblastoma cells. In fact, the pRb pathway is located downstream of EGFR, as EGFR regulates cyclin D expression and activation via the RAS/MAPK and the PI3K/Akt/mTOR pathway, respectively^48^. Yet there might be certain feedback mechanisms that are still unknown, for example via downstream E2F transcription factors, which regulate the expression of a plethora of genes^49^. However, if the link between *p16* knockout and EGFR inhibitor sensitivity would be through the canonical pRb pathway, one would expect that loss of p16 enhances CDK4/6 inhibitor sensitivity, which is in contrast with our findings. Interestingly, Fang et al. reported an inverse correlation between p27 and EGFR levels in human cancer tissues. They showed that depletion of p27 resulted in upregulation of EGFR via released suppression of the JNK/c-Jun pathway^50^. Moreover, inhibition of p27 phosphorylation at Ser^10^ led to p27 accumulation in the nucleus and enhanced erlotinib-mediated cytotoxicity^51^. Since the CDK4-cyclin D complex cannot bind p16 and p27 at the same time, more p27 might be directed to this complex when p16 is lost. Shuttling of p27 may result in released inhibition of the JNK/c-Jun pathway, which enhances EGFR expression. Although the link to EGFR was not investigated, p16 itself was also shown to bind JNK1/3 and block the activation of c-Jun^52^. Another possibility is that *p16* loss indirectly stimulates the release of growth factors, such as HBEGF and TGFβ1^53,54^, via an unknown mechanism. Future experiments looking further into the expression and localisation of EGFR, p27 and different ligands may help to understand the underlying mechanism.

We also observed significantly increased resistance to some ALK inhibitors in our *p16* knockouts. Since SY5Y cells carry an F1174L *ALK* mutation, these cells are not sensitive to every ALK inhibitor and increased resistance was only seen for those that are effective in cells with this specific mutation. Downregulation of ALK in the *p16* knockouts can explain this shift in sensitivity and further indicates a change of dependency on oncogenic pathways.

While we see a clear shift in phenotype when we knock out *p16*, the clinical implication of our findings requires further investigation. We showed that afatinib reduced tumour growth in SY5Y xenografts with *CDKN2A* knockout, but our results also indicate that combination strategies might be necessary to achieve remission. Future experiments will have to explore different drug combinations and confirm our results in other mouse models. The SY5Y cell line is interesting due to its SKNSH-derived clonal origin and its ability to transdifferentiate between a noradrenergic and a mesenchymal state^55^. Future experiments will have to elucidate the potential correlation between *p16/CDKN2A* and a phenotypic switch and investigate whether a mesenchymal transition occurs upon *p16* knockout in other neuroblastoma models. The tumour microenvironment plays an important role when it comes to mesenchymal transitions^56^, raising the question whether *p16* loss in patients induces the same changes as *in vitro*. Nonetheless, our findings fit with other observations that 1) homozygous loss of *CDKN2A* occurs predominantly in relapsed neuroblastoma^15^, 2) mesenchymal cells are less differentiated and enriched in relapse tumours^40,57^, and 3) the switch to a mesenchymal phenotype contributes to the development of relapses^58^. Similar to *RAS* mutations, *p16/CDKN2A* inactivation may be a way to induce neuronal dedifferentiation, activate stem cell characteristics and thereby contribute to therapeutic resistance and relapse. Overall, our study shows that *p16*, but not *p14*, knockout induces large transcriptomic changes in SY5Y neuroblastoma cells, as well as increased sensitivity to EGFR inhibitors. Our study paves that way for further research that aims at targeting *CDKN2A* deletions in neuroblastoma and potentially other (paediatric) cancer types.

## ACKNOWLEDGEMENTS

We thank the MCCA Intervention Unit of the Netherlands Cancer Institute for performing the *in vivo* experiments.

## REFERENCES

1. Campbell, P. J. et al. Pan-cancer analysis of whole genomes. Nature 578, 82–93 (2020).

2. Ma, X. et al. Pan-cancer genome and transcriptome analyses of 1,699 paediatric leukaemias and solid tumours. Nature 555, 371–376 (2018).

3. Sharpless, N. E. INK4a/ARF: A multifunctional tumor suppressor locus. Mutat. Res. -Fundam. Mol. Mech. Mutagen. 576, 22–38 (2005).

4. Li, Y., Nichols, M. A., Shay, J. W. & Xiong, Y. Transcriptional Repression of the D-Type Cyclin-dependent Kinase Inhibitor pl6 by the Retinoblastoma Susceptibility Gene Product pRb. Cancer Res. 54, 6078–6082 (1994).

5. Stott, F. J. et al. The alternative product from the human CDKN2A locus, p14(ARF), participates in a regulatory feedback loop with p53 and MDM2. EMBO J. 17, 5001–5014 (1998).

6. Udayakumar, T., Shareef, M. M., Diaz, D. A., Ahmed, M. M. & Pollack, A. The E2F1/Rb and p53/MDM2 pathways in DNA repair and apoptosis: Understanding the crosstalk to develop novel strategies for prostate cancer radiotherapy. Semin. Radiat. Oncol. 20, 258–266 (2010).

7. Carén, H. et al. High-resolution array copy number analyses for detection of deletion, gain, amplification and copy-neutral LOH in primary neuroblastoma tumors: Four cases of homozygous deletions of the CDKN2A gene. BMC Genomics 9, (2008).

8. Dreidax, D. et al. Low p14ARF expression in neuroblastoma cells is associated with repressed histone mark status, and enforced expression induces growth arrest and apoptosis. Hum. Mol. Genet. 22, 1735–1745 (2013).

9. Brodeur, G. M. Neuroblastoma: Biological insights into a clinical enigma. Nat. Rev. Cancer 3, 203–216 (2003).

10. Maris, J. M. Recent advances in neuroblastoma. N. Engl. J. Med. 362, 2202 (2010).

11. Johnsen, J. I., Dyberg, C. & Wickström, M. Neuroblastoma—A neural crest derived embryonal malignancy. Frontiers in Molecular Neuroscience vol. 12 9 (2019).

12. Pinto, N. R. et al. Advances in risk classification and treatment strategies for neuroblastoma. J. Clin. Oncol. 33, 3008–3017 (2015).

13. Tas, M. L. et al. Neuroblastoma between 1990 and 2014 in the Netherlands: Increased incidence and improved survival of high-risk neuroblastoma. Eur. J. Cancer 124, 47–55 (2020).

14. Moreno, L. et al. Outcome of children with relapsed or refractory neuroblastoma: A meta-analysis of ITCC/SIOPEN European phase II clinical trials. Pediatr. Blood Cancer 64, 25–31 (2017).

15. Eleveld, T. F. et al. Relapsed neuroblastomas show frequent RAS-MAPK pathway mutations. Nat. Genet. 47, 864–871 (2015).

16. Padovan-Merhar, O. M. et al. Enrichment of Targetable Mutations in the Relapsed Neuroblastoma Genome. PLOS Genet. 12, e1006501 (2016).

17. Takita, J. et al. Deletion Map of Chromosome 9 and p16 (CDKN2A) Gene Alterations in Neuroblastoma. Cancer Res. 57, (1997).

18. Marshall, B., Isidro, G., Martins, A. G. & Boavida, M. G. Loss of heterozygosity at chromosome 9p21 in primary neuroblastomas: Evidence for two deleted regions. Cancer Genet. Cytogenet. 96, 134–139 (1997).

19. Carr-Wilkinson, J. et al. High frequency of p53/MDM2/p14ARF pathway abnormalities in relapsed neuroblastoma. Clin. Cancer Res. 16, 1108–1118 (2010).

20. Schubert, N. A. et al. Combined targeting of the p53 and pRb pathway in neuroblastoma does not lead to synergistic responses. Eur. J. Cancer 142, 1–9 (2021).

21. Kamijo, T., Bodner, S., Van De Kamp, E., Randle, D. H. & Sherr, C. J. Tumor spectrum in ARF-deficient mice. Cancer Res. 59, 2217–2222 (1999).

22. Kamijo, T. et al. Tumor suppression at the mouse INK4a locus mediated by the alternative reading frame product p19(ARF). Cell 91, 649–659 (1997).

23. Krimpenfort, P., Quon, K. C., Mooi, W. J., Loonstra, A. & Berns, A. Loss of p16Ink4a confers susceptibility to metastatic melanoma in mice. Nature 413, 83–86 (2001).

24. Sharpless, N. E. et al. Loss of p16Ink4a with retention of p19 predisposes mice to tumorigenesis. Nature 413, 86–91 (2001).

25. Shalem, O. et al. Genome-scale CRISPR-Cas9 knockout screening in human cells. Science (80-.). 343, 84–87 (2014).

26. Sanjana, N. E., Shalem, O. & Zhang, F. Improved vectors and genome-wide libraries for CRISPR screening. Nature Methods vol. 11 783–784 (2014).

27. Brinkman, E. K., Chen, T., Amendola, M. & Van Steensel, B. Easy quantitative assessment of genome editing by sequence trace decomposition. Nucleic Acids Res. 42, e168–e168 (2014).

28. Y, L., Gk, S. & W, S. The R package Rsubread is easier, faster, cheaper and better for alignment and quantification of RNA sequencing reads. Nucleic Acids Res. 47, (2019).

29. Leek, J. et al. sva: Surrogate Variable Analysis. R package version 3.40.0 (2021).

30. Mi, L., W, H. & S, A. Moderated estimation of fold change and dispersion for RNA-seq data with DESeq2. Genome Biol. 15, (2014).

31. Korotkevich, G. et al. Fast gene set enrichment analysis. bioRxiv 060012 (2021) doi:10.1101/060012.

32. Liberzon, A. et al. The Molecular Signatures Database Hallmark Gene Set Collection. Cell Syst. 1, 417–425 (2015).

33. Butler, A., Hoffman, P., Smibert, P., Papalexi, E. & Satija, R. Integrating single-cell transcriptomic data across different conditions, technologies, and species. Nat. Biotechnol. 2018 365 36, 411–420 (2018).

34. Jansky, S. et al. Single-cell transcriptomic analyses provide insights into the developmental origins of neuroblastoma. Nat. Genet. 1–11 (2021) doi:10.1038/s41588-021-00806-1.

35. Twentyman, P. R. & Luscombe, M. A study of some variables in a tetrazolium dye (MTT) based assay for cell growth and chemosensitivity. Br. J. Cancer 56, 279–285 (1987).

36. Kovalevich, J. & Langford, D. Considerations for the use of SH-SY5Y neuroblastoma cells in neurobiology. Methods Mol. Biol. 1078, 9–21 (2013).

37. Jakacki, R. I. et al. Pediatric Phase I and Pharmacokinetic Study of Erlotinib Followed by the Combination of Erlotinib and Temozolomide: A Children’s Oncology Group Phase I Consortium Study. https://doi.org/10.1200/JCO.2007.15.2306 26, 4921–4927 (2016).

38. Jp, T., H, A., Ry, H. & Ma, N. Epithelial-mesenchymal transitions in development and disease. Cell 139, 871–890 (2009).

39. Lamouille, S., Xu, J. & Derynck, R. Molecular mechanisms of epithelial-mesenchymal transition. Nat. Rev. Mol. Cell Biol. 15, 178–196 (2014).

40. Gartlgruber, M. et al. Super enhancers define regulatory subtypes and cell identity in neuroblastoma. Nat. Cancer 2, 114–128 (2021).

41. Al-Khalaf, H. H. & Aboussekhra, A. p16INK4A induces senescence and inhibits EMT through microRNA-141/ microRNA-146b-5p-dependent repression of AUF1. Mol. Carcinog. 56, 985–999 (2017).

42. Hecht, M., Papoutsi, M., Tran, H. D., Wilting, J. & Schweigerer, L. Hepatocyte Growth Factor/c-Met Signaling Promotes the Progression of Experimental Human Neuroblastomas. Cancer Res. 64, 6109–6118 (2004).

43. Lo, H. W. et al. Epidermal growth factor receptor cooperates with signal transducer and activator of transcription 3 to induce epithelial-mesenchymal transition in cancer cells via up-regulation of TWIST gene expression. Cancer Res. 67, 9066–9076 (2007).

44. Kim, J., Kong, J., Chang, H., Kim, H. & Kim, A. EGF induces epithelial-mesenchymal transition through phospho-Smad2/3-Snail signaling pathway in breast cancer cells. Oncotarget 7, 85021–85032 (2016).

45. Katz, M. et al. A reciprocal tensin-3-cten switch mediates EGF-driven mammary cell migration. Nat. Cell Biol. 9, 961–969 (2007).

46. Lemmon, M. A., Freed, D. M., Schlessinger, J. & Kiyatkin, A. The Dark Side of Cell Signaling: Positive Roles for Negative Regulators. Cell 164, 1172–1184 (2016).

47. Avraham, R. & Yarden, Y. Feedback regulation of EGFR signalling: Decision making by early and delayed loops. Nat. Rev. Mol. Cell Biol. 12, 104–117 (2011).

48. Wee, P. & Wang, Z. Epidermal growth factor receptor cell proliferation signaling pathways. Cancers (Basel). 9, (2017).

49. Bracken, A. P., Ciro, M., Cocito, A. & Helin, K. E2F target genes: unraveling the biology. doi:10.1016/j.tibs.2004.06.006.

50. Fang, Y. et al. A new tumour suppression mechanism by p27Kip1: EGFR down-regulation mediated by JNK/c-Jun pathway inhibition. Biochem. J. 463, 383–392 (2014).

51. Zhang, D. et al. Silencing kinase-interacting stathmin gene enhances erlotinib sensitivity by inhibiting ser10 p27 phosphorylation in epidermal growth factor receptor-expressing breast cancer. Mol. Cancer Ther. 9, 3090–3099 (2010).

52. Choi, B. Y. et al. The tumor suppressor p16 INK4a prevents cell transformation through inhibition of c-Jun phosphorylation and AP-1 activity. (2005) doi:10.1038/nsmb960.

53. Schneider, M. R. & Wolf, E. The epidermal growth factor receptor ligands at a glance. J. Cell. Physiol. 218, 460–466 (2009).

54. Zhao, Y. et al. TGF-β transactivates EGFR and facilitates breast cancer migration and invasion through canonical Smad3 and ERK/Sp1 signaling pathways. Mol. Oncol. 12, 305–321 (2018).

55. Thirant, C. et al. Interplay between intrinsic reprogramming potential and microenvironment controls 1 neuroblastoma cell plasticity and identity 2 3. bioRxiv 2021.01.07.425710 (2021) doi:10.1101/2021.01.07.425710.

56. Jung, H.-Y., Fattet, L. & Yang, J. Molecular Pathways Molecular Pathways: Linking Tumor Microenvironment to Epithelial-Mesenchymal Transition in Metastasis. Clin Cancer Res 21, 1–7 (2014).

57. Van Groningen, T. et al. Neuroblastoma is composed of two super-enhancer-associated differentiation states. Nat. Genet. 49, 1261–1266 (2017).

58. Mitra, A., Mishra, L. & Li, S. EMT, CTCs and CSCs in tumor relapse and drug-resistance. Oncotarget 6, 10697–10711 (2015).

