## Supplementary Figure 1 for "Loss of *p16INK4a* in neuroblastoma cells induces shift to an immature state with mesenchymal characteristics and increases sensitivity to EGFR inhibitors"

a)

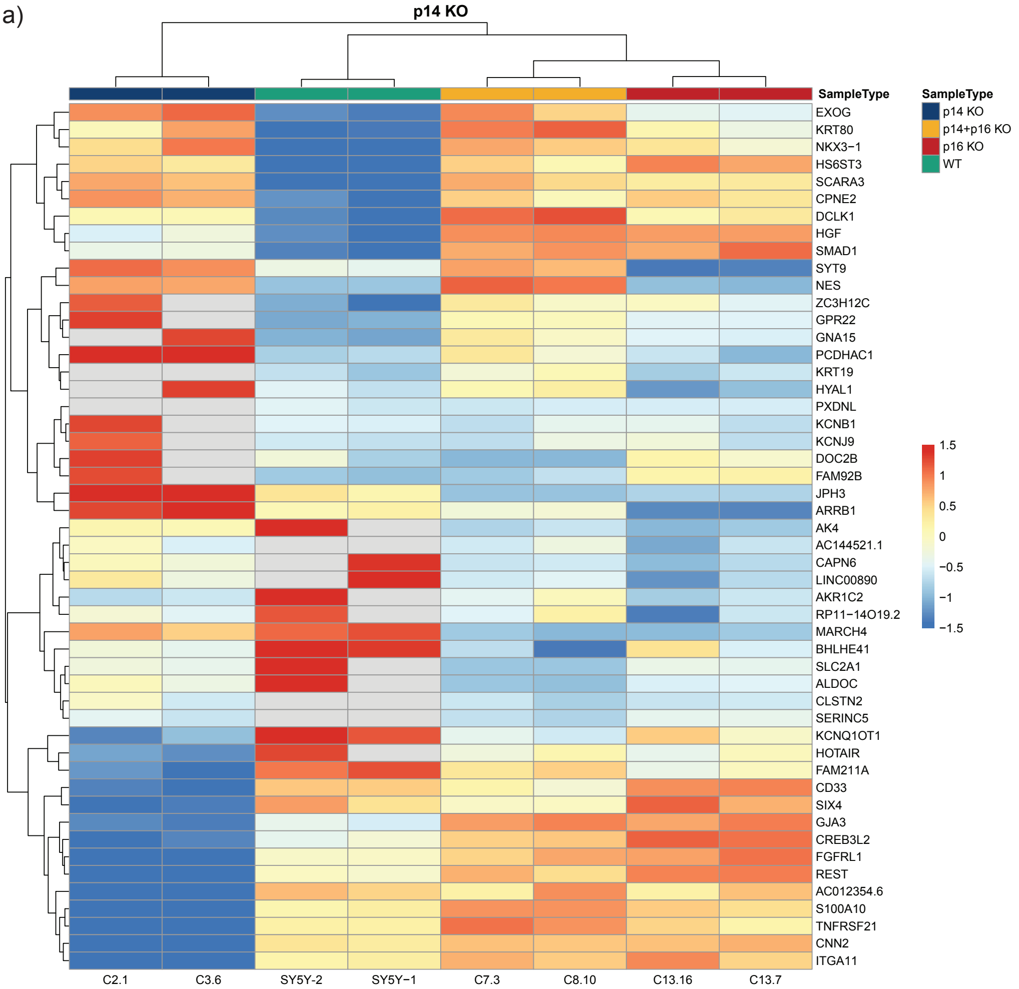


b)

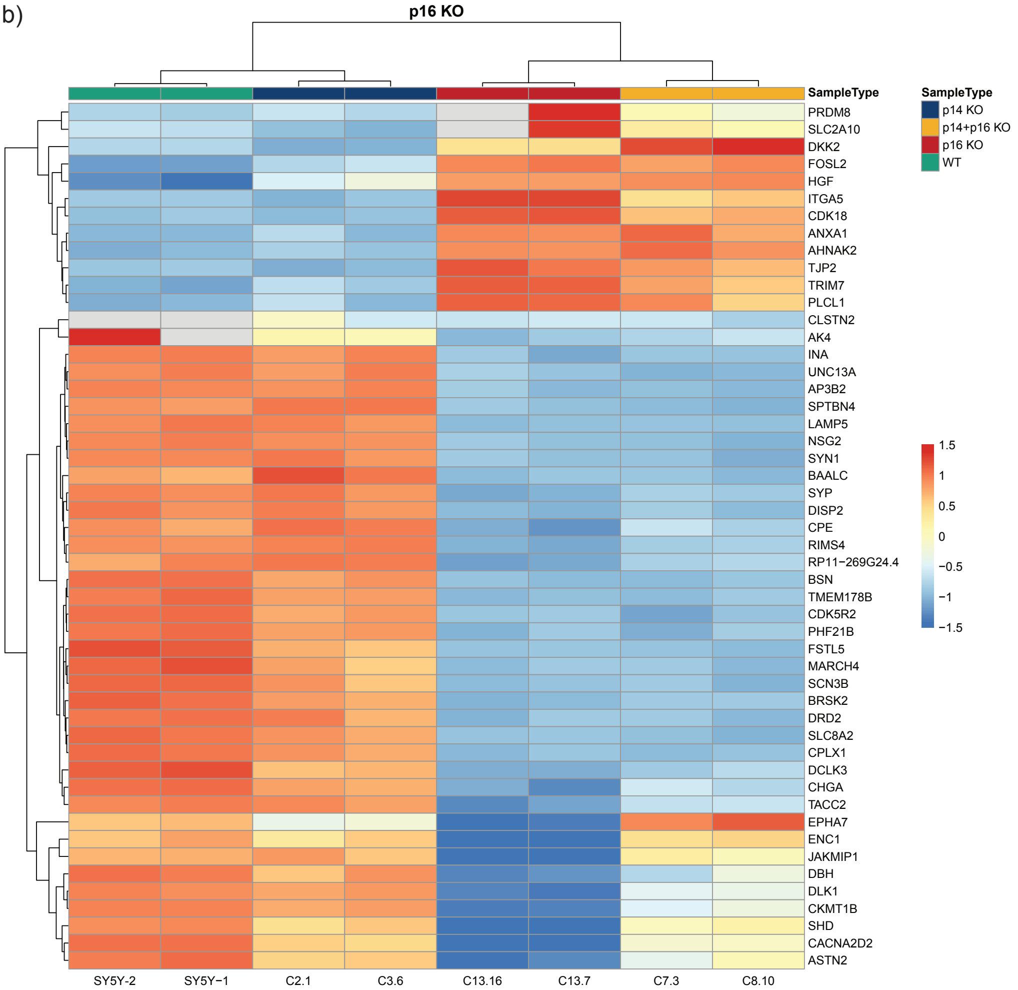


**Supplementary Figure 1** Gene expression of the different knockout cell lines. **a)** Heatmap showing the expression (transcripts per million) of the 50 most differentially expressed genes (*p14* knockout versus wildtype) with an absolute log fold change >1. **b)** Heatmap showing the expression (transcripts per million) of the 50 most differentially expressed genes (*p16* knockout versus wildtype) with an absolute log fold change >1. Genes are standardised row-wise.
