## Supplementary Figure 2 for "Loss of *p16INK4a* in neuroblastoma cells induces shift to an immature state with mesenchymal characteristics and increases sensitivity to EGFR inhibitors"

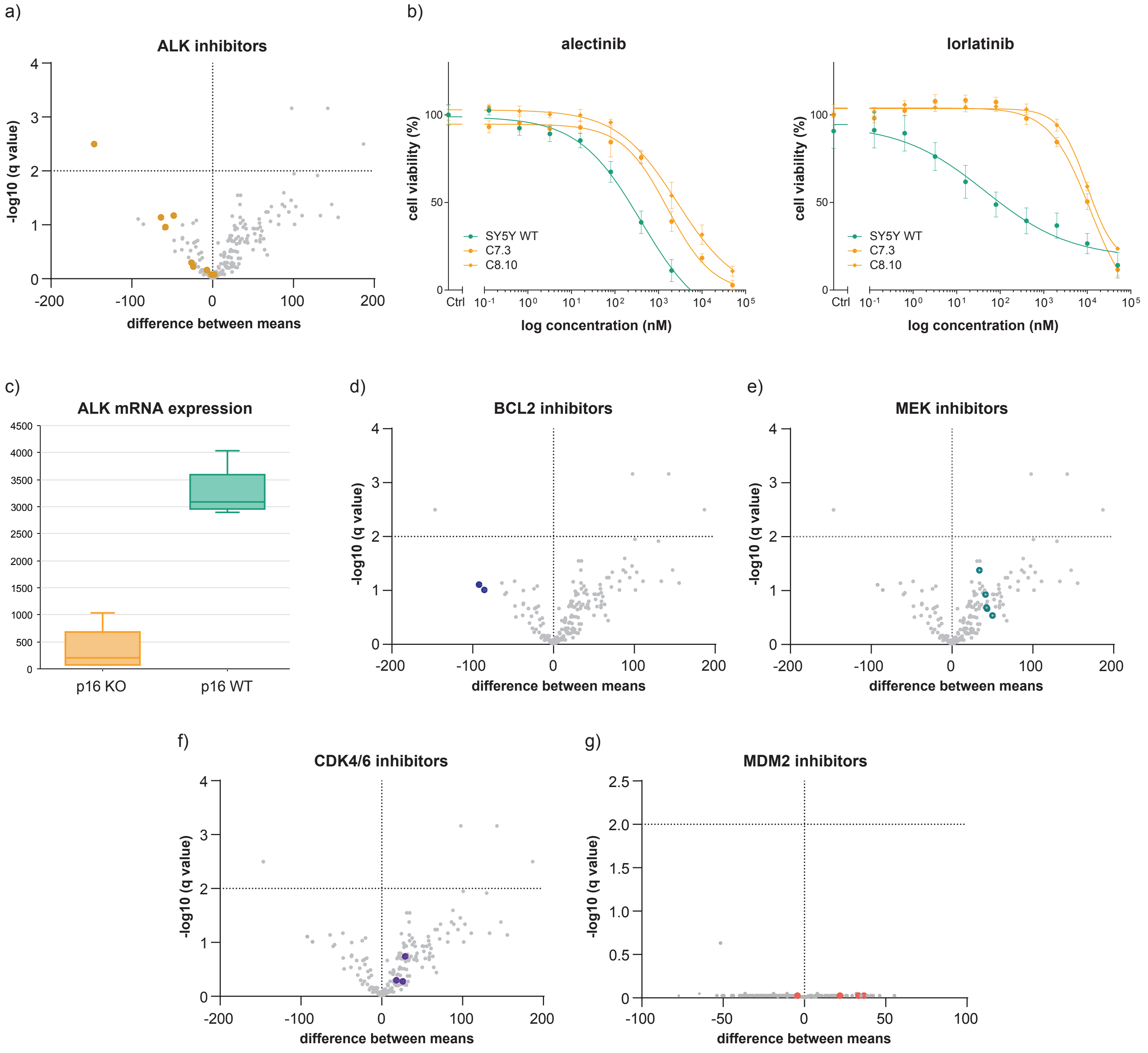


**Supplementary Figure 2** Sensitivities of the different knockout cell lines for ALK, BCL2, MEK, CDK4/6 and MDM2 inhibitors. **a)** Volcano plot comparing area under the curve (AUC) values of *p16* wildtype (SY5Y wildtype and *p14* knockout) with *p16* knockout (*p16* and *p14+p16* knockout) cell lines. All nine ALK inhibitors included in the library are highlighted. **b)** Validation of resistance to ALK inhibitors alectinib and lorlatinib in *p14+p16* knockout cells. Error bars indicate the SEM. **c)** Boxplot showing ALK mRNA expression in *p16* knockout and *p16* wildtype clonal cell lines. **d+e)** Volcano plot comparing AUC values for BCL2 inhibitors **(e)** and MEK inhibitors **(f)** of *p16* wildtype (SY5Y wildtype and *p14* knockout) with *p16* knockout (*p16* and *p14+p16* knockout) cell lines. BCL2 (blue) and MEK (green) inhibitors included in the library are highlighted. **f+g)** Volcano plot comparing AUC values of *p16* wildtype (SY5Y wildtype and *p14* knockout) with *p16* knockout (*p16* and *p14+p16* knockout) cell lines **(f)** or *p14* wildtype (SY5Y wildtype and *p16* knockout) with *p14* knockout (*p14* and *p14+p16* knockout) cell lines **(g)**. CDK4/6 (purple) or MDM2 (coral) inhibitors included in the library are highlighted.
